# Single Session Real-time fMRI Neurofeedback has a Lasting Impact on Cognitive Behavioral Therapy Strategies

**DOI:** 10.1101/258095

**Authors:** Katherine E. MacDuffie, Jeff MacInnes, Kathryn C. Dickerson, Kari M. Eddington, Timothy J. Strauman, R. Alison Adcock

**Affiliations:** Department of Speech and Hearing Sciences, University of Washington, Seattle, WA; Department of Psychology, University of Washington, Seattle, WA; Department of Psychiatry and Behavioral Sciences, Duke University, Durham, NC.; Department of Psychology, University of North Carolina at Greensboro, Greensboro, NC.; Department of Psychology and Neuroscience, Duke University, Durham, NC.; Department of Neurobiology, Duke University, Durham, NC.

**Keywords:** real-time fMRI, neurofeedback, cognitive-behavioral therapy, metacognition, personalized therapy

## Abstract

To benefit from cognitive behavioral therapy (CBT), individuals must not only learn new skills but also strategically implement them outside the session. Here, we tested a novel technique for personalizing CBT skills and facilitating their generalization to daily life. We hypothesized that showing participants the impact of specific CBT strategies on their own brain function using real-time functional magnetic imaging (rt-fMRI) neurofeedback would increase their metacognitive awareness, help them identify effective strategies, and motivate real-world use. In a within-subjects design, participants who had completed a clinical trial of a standardized course of CBT created a personal repertoire of negative autobiographical stimuli and mood regulation strategies. From each participant’s repertoire, a set of experimental and control strategies were identified; only experimental strategies were practiced in the scanner. During the rt-fMRI neurofeedback session, participants used negative stimuli and strategies from their repertoire to manipulate activation in the anterior cingulate cortex, a region implicated in emotional distress. The primary outcome measures were changes in participant ratings of strategy difficulty, efficacy, and frequency of use. As predicted, ratings for unscanned control strategies were stable across observations, whereas ratings for experimental strategies changed after neurofeedback. At follow-up one month after the session, efficacy and frequency ratings for scanned strategies were predicted by neurofeedback during the rt-fMRI session. These results suggest that rt-fMRI neurofeedback created a salient and durable learning experience for patients, extending beyond the clinic to guide and motivate CBT skill use weeks later. This metacognitive approach to neurofeedback offers a promising model for increasing clinical benefits from cognitive-behavioral therapy by personalizing skills and facilitating generalization.

## Introduction

Efficacious psychotherapies convey clinical benefit in part through learning, often relying on the appropriate application of new skills (Kazdin, 2007). For example, individuals with a disorder like depression may benefit from developing the ability to challenge (or change) negative thoughts, increase positive behaviors, and cope with distressing emotions. While these skills can be practiced during a therapy session, the best outcomes are obtained when individuals successfully incorporate new skills into their daily routines (Hundt, Mignogna, Underhill, & Cully, 2013). However, generalization to life outside of the therapy session remains one of the critical challenges of psychotherapy. As dissemination of evidence-based therapies occurs on a broader scale (Karlin & Cross, 2014), methods to increase successful generalization of therapeutic learning and optimize real-world skill use are needed to improve clinical outcomes.

Several challenges to generalization may be characterized as metacognitive: a patient may have been told that a new skill can be helpful, but without having experienced benefits directly, he or she may not be motivated to use it; we refer to this problem as one of *credibility*. Alternatively, a patient may have tried a strategy previously and found it to be effective, but if the skill has not yet become a part of his or her daily behavioral repertoire, the cue to implement it depends on its retrievability from memory. We refer to this problem as one of *availability*—if a given skill is not available in memory to be retrieved and used when needed, then it is unlikely to be used outside of session. Critically, since skills must be practiced outside of session to be adequately reinforced, these metacognitive barriers may be an overriding constraint on generalization and progress. Thus, the state of an individual’s metacognitive awareness and knowledge about therapeutic techniques may either limit or permit gains from psychotherapy.

An additional challenge concerns how to apply the skills gained in psychotherapy flexibly in response to diverse real-world scenarios. Most learning-based therapies include a rich repertoire of skills and concepts that must be adapted across scenarios and individuals. Selection of a specific strategy for use in daily life is not trivial, and monitoring the impact of a strategy on one’s own symptoms is a complex, cognitively-demanding problem. The ability to efficiently compare and contrast the effects of specific strategies on one’s own symptoms and psychobiology is therefore another critical constraint on clinical benefit. Improving patients’ ability to identify the specific skills that are the most effective for them should support the development of a personalized repertoire of skills and optimize real-world skill use.

### Increasing credibility and availability of therapy skills

The notion that psychotherapy can change the brain is intuitive to neuroscientists; neuroscience conceptualizes therapy as a learning experience that confers long-term benefits precisely via alteration of brain structure and function. However, members of the general public are likely to believe that psychotherapy alone is insufficient for treating “biological” mental disorders (Deacon & Baird, 2009). We aimed to counter this assumption by harnessing the persuasive power of neuroscience information, which appeals to and is considered highly credible by lay audiences (Fernandez-Duque, Evans, Christian, & Hodges, 2015; McCabe & Castel, 2008; Weisberg, Keil, Goodstein, Rawson, & Gray, 2008). We hypothesized that a live demonstration of *therapy skills changing brain activity* would be particularly effective for increasing the credibility of skills learned in therapy. We created such a demonstration using real-time functional magnetic resonance imaging (rt-fMRI) neurofeedback.

In addition to improving credibility, a neurofeedback demonstration of brain activity changing in response to therapy skills is also likely to be salient and memorable for participants, thus increasing the availability of practiced skills. Rt-fMRI neurofeedback is a relatively new technology and one that few people have had the opportunity to experience directly (Sulzer, Haller, Scharnowski, & Weiskopf, 2013). Therefore, we hypothesized that the experience would be novel, and thus more likely to be remembered compared to a more routine laboratory or clinic procedure.

### Personalized feedback to aid skill selection

Rt-fMRI neurofeedback is an emerging technology with significant advantages over feedback modalities like EEG or psychophysiology; it offers increased access to subcortical areas and, importantly, anatomical and temporal specificity (Linden et al., 2012; Sulzer et al., 2013; Young et al., 2014). Visual feedback depicting activation in a specific structure can be displayed in real-time, and used to train brain activation or to demonstrate volitional control of brain activity. In the current study, we aimed to show participants the efficacy of specific skills they had learned in therapy to change brain activity, using an idiographic approach that offered neurofeedback corresponding to a personalized repertoire of strategies. This approach aimed not only to enhance general credibility and motivate real-world skill use, but also to inform evaluation of specific skills to improve selection.

### A novel approach to rt-fMRI neurofeedback

The current study aimed to test whether a single rt-fMRI session could enhance metacognitive awareness of the neural changes associated with use of existing skills for regulating negative emotions. We built upon rt-fMRI interventions for depression where participants were trained to upregulate brain activation responsive to negative emotions (Linden et al., 2012; Young et al., 2014). Unlike prior studies, however, we did not attempt to teach a new regulation skill to our participants. Here, the scan session consisted entirely of a “test” phase, in which participants could draw their own conclusions about associations between neural activation and previously-learned mood regulation skills. To achieve this, we recruited participants who had prior experience with CBT and a presumably well-learned set of skills that could be tested inside the MRI scanner.

Our approaches for selecting regions of interest (ROI) and stimuli for the scan session were idiographic, prioritizing ecological validity and individual differences over standardization. An ROI search was conducted within the anatomical boundaries of the anterior cingulate cortex (ACC) because activation in this region has been shown to correlate with subjective distress and has favorable signal-to-noise ratio for providing real-time neurofeedback (Hamilton, Glover, Hsu, Johnson, & Gotlib, 2011). We selected a unique region of interest (ROI) for each participant to maximize our chances of providing a strong neurofeedback signal to each individual. The stimuli for the scan session were also personalized. Participants reported previously-learned strategies that were effective for controlling mood. Those that were purely cognitive (e.g., checking the facts) were compatible with fMRI and thus could be used in the scanner. Those that were behavioral (e.g., calling a friend), or physical (e.g., breath control) could not be used in the scanner, and were analyzed as within-subject control strategies.

To our knowledge, this is the first rt-fMRI study to use a metacognitive rather than feedback-training approach, and the first time that fMRI neurofeedback has been used to support generalization of previously-learned mood regulation strategies. We predicted significant correspondence between the neurofeedback received and outcomes related to skill generalization: subjective *efficacy* (a measure of credibility) and *frequency* (a measure of availability) of specific strategy use, one month later. We hypothesized that only those strategies that successfully decreased ACC activation would be rated as more effective and more frequently used following the neurofeedback experience. Similarly, we predicted that, across participants, those who had more success at decreasing ACC activation with their strategies would endorse higher efficacy and frequency of strategy use at follow-up.

## Methods and Materials

### Participants

We recruited from a group of 24 participants who had prior diagnoses of depression and had previously completed a standardized course of CBT in a randomized clinical trial (Eddington et al., 2015). We chose this recruitment approach to minimize treatment heterogeneity, as all participants had received equivalent doses of evidence-based treatment delivered in the same clinical setting. Recruiting from this advantageous sample, however, limited our potential sample size; thus, we elected to deliver the active intervention to all participants and utilize a within-subject control.

Potential participants were excluded if they were deemed ineligible for the MRI scan (due to medical/safety concerns; *n* = 3), reported active suicidal ideation and a history of previous attempts (*n* = 1), or were not interested in participating (*n* = 7). A final sample of 13 participants (11 F; mean age = 44) took part in the scan session.

Participants were recruited for their prior therapy experience, rather than diagnostic status, and served as their own controls. Thus, a full diagnostic assessment was not performed. Before enrolling in the initial CBT treatment trial, a majority of the participants (8/13) had been naïve to therapy and all were medication-naïve. Since completing the trial, three had pursued additional therapy and four had started taking antidepressant medications. On average, 29 months had elapsed since CBT treatment completion (range: 19-39 months); thus, we expected and observed variability in mood symptoms. The sample was characterized by mild depressive symptoms at enrollment, with a mean PHQ-9 score of 8.2 (range: 1-18), and mean BDI-II score of 16 (range: 0-30).

### Experimental timeline and strategy ratings

The study took place over a 6-week span (Fig 1). The Pre-Scan Interview (week 1) and Scan Session (week 2) were both completed in person. The Follow-Up assessment (week 6) was completed remotely using the Qualtrics survey platform. Screening measures were administered at each timepoint for symptom and safety monitoring (i.e., assessing for suicidal ideation or acute distress), but were not primary outcome measures and are not relevant to the current report. All study procedures met ethical standards and were approved by the Duke Medicine Institutional Review Board.

**Fig 1.**
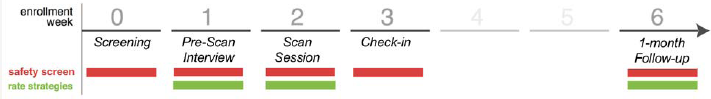
Timeline of Study Events. Screening measures were administered at each timepoint for symptom and safety monitoring. At Week 1, the Pre-Scan Interview catalogued and codified each individual participant’s repertoire of negative memories (or worries) and of CBT strategies for regulating negative emotion. At Week 1 (Baseline) and Week 6 (Follow-Up), all strategies were rated for *frequency, efficacy*, and *difficulty* of use. At Week 2, immediately following the rt-fMRI neurofeedback task runs (Scan Session), all scanned strategies were rated for efficacy and difficulty.

Strategy ratings were collected at the Pre-Scan Interview (hereafter *Baseline*), during the *Scan Session* (immediately after participants finished the real-time task runs), and at the 1-month *Follow-Up* (Fig 1). Participants used 7-point Likert-type scales to rate each strategy in terms of efficacy, difficulty of use, and frequency of use. Our primary hypotheses concerned strategy *efficacy*, and *frequency* of use ratings, which we expected to change in response to neurofeedback. We also collected ratings of *difficulty*, which we did not expect to change in response to neurofeedback.

### Pre-scan interview session

A two-hour in-person Pre-Scan Interview was conducted one week prior to the Scan Session. Participants reviewed their therapy experiences and listed 24 negative memories or worries that could be used to trigger negative emotions in the scanner. Each memory or worry was rated for emotionality and level of detail, and identified with a short phrase to be shown in the scanner (e.g., “pool incident” or “Mom’s overdue medical bills.”)

Participants also identified 8 strategies, learned in therapy or elsewhere, that they believed effective for regulating their mood. All strategies were rated for efficacy, difficulty, and frequency of use. Each strategy was identified with a short phrase to be shown in the scanner (e.g., “examine the evidence”).

Each participant’s repertoire of strategies was then divided to supply the *experimental* (for use during the MRI session) and *control* conditions. The assignment of strategies was not fully random, because some of the strategies listed were incompatible with the MRI environment (e.g., deep breathing) and were necessarily assigned to the control condition. This stratification nevertheless allowed us to compare the stability of ratings across time for strategies used or not used during the scan session.

### Scan session

The scan session took place one week after the Pre-Scan Interview. Participants remained in the scanner for 1.5-2 hours during collection of structural images, functional localizers, resting state scans, and up to 5 real-time neurofeedback runs.

For each participant, their individual experimental strategies (as identified during the Pre-Scan Interview) were pseudo-randomly paired with the participant’s 24 negative memories/worries, such that each strategy appeared an equal number of times. The memory-strategy pairs were randomized for use during the localizer and real-time neurofeedback runs.

#### Localizer task

The localizer task was comprised of 4 trials. Each trial cycled through three phases— Count, Memory, Strategy—with each phase lasting 30 seconds (Fig 2). During the *Count* phase, participants counted backwards from a starting value using a specified increment (e.g., 300 by 4). Task instructions for the count phase minimized the potential stress effects of mental subtraction by emphasizing the absence of a target value or penalty for errors. During the *Memory* phase, participants viewed an idiographic <u>memory/worry</u> cue and were instructed to elaborate on the emotions associated with the memory or worry. During the *Strategy* phase, participants viewed an idiographic <u>strategy</u> cue and were instructed to employ that specific strategy to regulate their mood.

**Fig 2.**
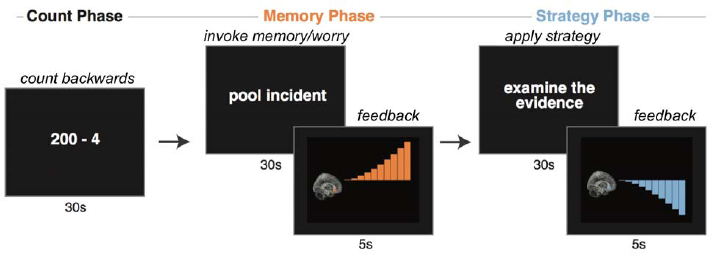
Rt-fMRI Neurofeedback Task Design. The rt-fMRI neurofeedback trial structure is illustrated here. The localizer task was identical except it did not include feedback. Activation in the individualized region of interest (ROI) was presented as neurofeedback following negative memory retrieval or worry (Memory Phase) and regulation (Strategy Phase). Successive trials rotated through each participant’s repertoire of memories and strategies, using the personalized codes generated in the Pre-Scan Interview.

#### Real-time neurofeedback task

The real-time neurofeedback task was identical in structure and timing to the localizer task, except that following each *Memory* and *Strategy* phase, participants viewed a brief (5 s) neurofeedback display indicating fMRI activation during the preceding phase (See Fig 2). Participants completed 5 real-time runs, consisting of 4 trials each (˜8 min/run). To minimize potential distress, after each run, participants were offered the choice to continue or to end the session.

### Imaging parameters

#### fMRI acquisition

Functional runs were collected on a 3.0T GE MR750 scanner using an echo-planar sequence with the parameters: TR, 1 s; TE, 28 ms; flip angle, 90°; voxel size, 3x3x3.8 mm; 18 oblique axial slices, parallel to the anterior commissure - posterior commissure axis. The first 8 volumes of each run were discarded to permit stabilization of the net magnetization. Fast spoiled gradient echo high-resolution whole-volume T1-weighted images (voxel size, 1x1x1 mm) were acquired.

#### Real-time fMRI neurofeedback

The five task runs included neurofeedback after each *Memory* and *Strategy* phase, calculated from a participant-specific ACC ROI (see *ROI Definition*). Throughout the scan a dedicated real-time analysis computer retrieved reconstructed imaging data, calculated ROI averages, and made those results available to the neurofeedback presentation computer. The neurofeedback presentation consisted of an ascending or descending series of animated signal bars that conveyed the magnitude of fMRI brain activation change in each phase (See Fig 2).

For the *Memory* phase, participants were instructed to expect increases in brain activation. The neurofeedback value was calculated as the average ROI signal during the last 15 seconds of the *Memory* phase minus the average ROI signal during the last 15 seconds of the preceding *Count* phase (baseline). The feedback range was normalized to [0 to 1] to present relative changes, with any negative values truncated at 0.

For the *Strategy* phase, participants were instructed to expect decreases in brain activation. The neurofeedback value was calculated as the average ROI signal during the last 15 seconds of the *Strategy* phase minus the average ROI signal during the last 15 seconds of the preceding *Memory* phase. The feedback range was normalized to [-1 to 0] to present relative changes, with any positive values truncated at 0.

The analytic focus for the present report was the neurofeedback provided to participants, showing the efficacy of strategies (specific and collective) at reducing activation in the Strategy phase, relative to the Memory phase.

#### ROI definition

Participant-specific ROIs were defined using the localizer task. We first ran a GLM modeling activation for the Memory Phase > [Strategy + Count Phases]. The resulting whole brain z-stat maps were then smoothed at 4 mm FWHM and masked to isolate activated voxels within an anatomically-defined ACC ROI (union of Brodmann Areas 24, 25 and 32; Maldjian, Laurienti, Kraft, & Burdette, 2003). Voxels within this ROI were segregated into clusters based on an activation threshold set at the 95^th^ percentile of all voxel activation. We selected the largest resultant cluster, identified the peak voxel, and constructed a 7-mm spherical ROI centered on this voxel for neurofeedback. For one participant, the peak voxel was from a whole-brain search—and not restricted to anatomical ACC—due to a registration error. Neurofeedback values for this participant did not differ from participants whose ROIs fell within the typical search area. For the majority of participants (8/13), the neurofeedback ROI was located in the paracingulate gyrus (Harvard-Oxford anatomical atlas; Fig 3).

**Fig 3.**
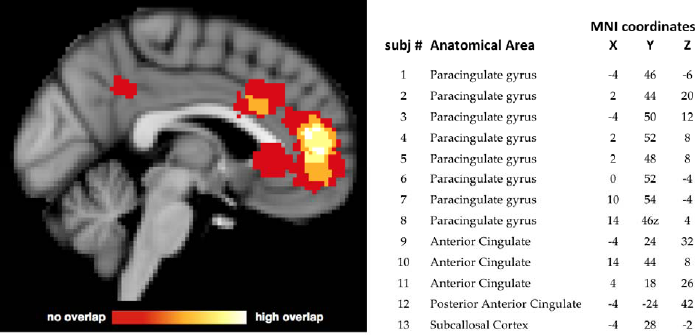
Location of Subject-Specific MNI-space ROIs. ROI selection used activation in the individual localizer runs intersected with anatomical priors. Anatomical boundaries reflected prior research implicating anterior cingulate cortex (ACC) in subjective distress. The center coordinate of each subject-specific ROI is listed along with anatomical label derived from the Harvard-Oxford structural atlas.

### Data analysis

Our analysis approach focused on whether the neurofeedback experience in the rt-fMRI session predicted changes in strategy efficacy and frequency ratings. 4 participants completed fewer than 20 neurofeedback trials; however, there were no associations between the number of neurofeedback trials completed and any of the outcome measures; all *p*s > .2). Pearson correlation coefficients were calculated to assess the relationship between trial-by-trial neurofeedback values and subjective strategy ratings collected at three observation intervals: baseline, during the scan session, and at 1-month follow-up. To account for the nested structure of the data, multi-level model analyses included observation interval at the lower level, and participant at the upper level. With a sample size of 13, statistical power is limited to detect effects in such a model (Maas & Hox, 2005). We therefore modeled only fixed effects, as our number of observations per participant (8 strategies, maximum 4 per condition) was insufficient to model individual differences in slopes.

## Results

We hypothesized that observing real-time fMRI neurofeedback depicting the ability of specific strategies to decrease brain activation generated by negative memories or worries would impact participants’ awareness of the *efficacy* of those strategies, and increase the *frequency* of their use in daily life, but not their perceived *difficulty*. To test these hypotheses, we looked for correspondence between presented neurofeedback and strategy ratings before versus during the scan session and at 1-month follow-up. We furthermore looked for changes in ratings (between baseline and follow-up) for strategies used in the scanner (*experimental*) compared to those that were not used (*control*). For one participant, follow-up strategy ratings were not obtained due to a technical error, leaving a sample size of 12 participants for the follow-up analyses reported below.

### Experimental vs. control strategies

Experimental and control strategies did not differ on average ratings of frequency, efficacy or difficulty, either at baseline or at follow-up (univariate tests in repeated measures ANOVA, all *p*s > .2). Thus, while experimental strategies may have differed qualitatively from control strategies, there was no significant quantitative difference in their average effectiveness, difficulty, or frequency of use.

Within-subject ratings of control strategies were significantly correlated at baseline and at follow-up (efficacy: *r* = .60, *p* = .04; frequency: *r* = .70, *p* = .01; difficulty: *r* = .57, *p* = .05), suggesting no significant change in the control strategies following completion of the fMRI session. In contrast, for experimental strategies, ratings at follow-up were not correlated with ratings at baseline (efficacy: *r* = .13, *p* = .68; frequency: *r* = .41, *p* = .18; difficulty: *r* = .10, *p* = .76).

### Correlations between strategy neurofeedback and strategy ratings

To examine whether the subjective experience of strategy efficacy during the scan session was related to the trial-by-trial presentation of neurofeedback, the feedback and ratings were collapsed across all subjects, yielding a total of 50 strategies. Individual strategies that were associated with a stronger neurofeedback signal (defined as decrease for Strategy relative to Memory in the ROI) were rated as more effective, *r* = -.31, *p* = .03, and easier to use, *r* = -.35, *p* = .01, on the day of the scan (Fig 4). At follow-up four weeks after the scan, strategies that were associated with a stronger feedback signal were rated as more effective, *r* = -.33, *p* = .03, and more frequently used (at trend level), *r* = -.26, *p* = .08. This supported our hypothesis that strategies for which neurofeedback demonstrated decreased ROI activation during the scan would be selectively associated with greater self-reported credibility and availability. Neurofeedback was not associated with follow-up difficulty ratings, however, *r* = -.12, *p* = .4, suggesting that strategies associated with stronger neurofeedback, while easier to use in the scanner, were not easier to use in daily life.

**Fig 4.**
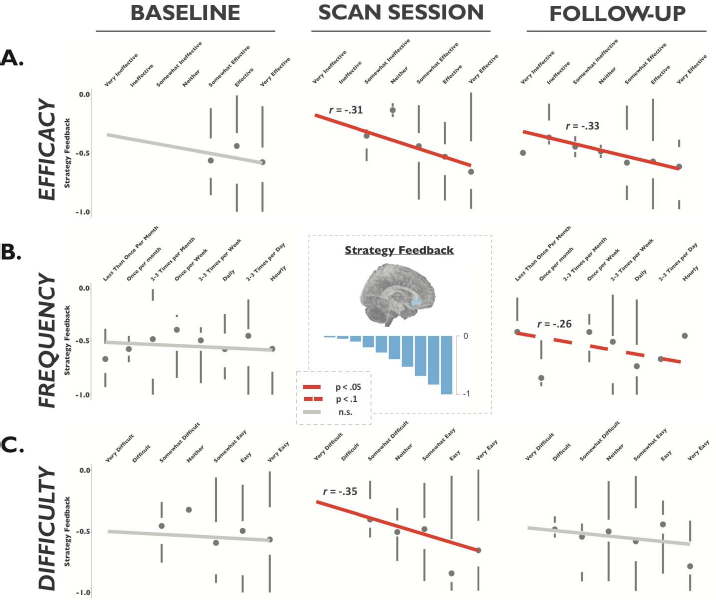
Strategy-Neurofeedback Correlations. Note that negative neurofeedback values indicate a reduction in activation from Memory to Strategy Phase. Neurofeedback was not associated with baseline ratings on any measure (first column). **A) Efficacy ratings**. Strategies associated with stronger neurofeedback (i.e., a larger reduction) were rated as significantly more effective during the scan session and at 1-month follow-up. **B) Frequency ratings**. Strategies associated with stronger neurofeedback showed a trend towards being rated as more frequently used at 1-month follow-up. **C) Difficulty ratings**. Strategies associated with stronger neurofeedback were rated as easier to use on the day of the scan session, but not at 1-month follow-up.

Critical to a conservative test of our hypothesis, we observed that pre-scan baseline ratings of frequency and efficacy for individual strategies were not predictive of feedback values received during the scan (Fig 4). For efficacy, the correlation of baseline ratings with strategy feedback was non-significant, *r* = -.12, *p* = .41, and statistically different from the correlation of strategy feedback with 1-month follow-up efficacy ratings, Steiger’s *Z* = 2.11, *p* = .03. For frequency of use, the correlation of baseline ratings with strategy feedback was non-significant, *r* = -.08, *p* = .60, and different from the coefficient with 1-month follow-up frequency ratings at trend-level, Steiger’s *Z* = 1.74, *p* = .08. For difficulty, the correlation of baseline ratings with strategy feedback was non-significant, *r* = -.06, *p* = .65, and not statistically different from the correlation of strategy feedback with 1-month follow-up difficulty ratings, Steiger’s *Z* = .331, *p* = .74. Thus, for scanned strategies, baseline ratings did not predict real-time fMRI neurofeedback values, but neurofeedback values did predict efficacy and frequency ratings four weeks after scanning.

### Multi-level model

Taking into account the nested structure of the data, the results of the multi-level model predicting efficacy ratings from neurofeedback confirmed the findings from the correlational analysis described above (Table 1). At the between-subjects level, subjects who received stronger neurofeedback values rated their strategies as more effective at 1-month follow-up, β1 = -1.94, *t* = 2.84, *p* < .01, but not at baseline, β0 = -.38, *t* = -.58, *p* = .56. The difference between the baseline and follow-up coefficients was significant at trend-level, *t* = -1.74, *p* = .08. Similarly, at the within-subject (strategy) level, strategies that were associated with higher neurofeedback values in the scanner were rated as significantly more effective at 1-month follow-up, β1 = -1.70, *t* = 2.53, *p* = .01, but not at baseline. β0 = -.06, *t* = -.1, *p* = .92. The difference between the baseline and follow-up coefficients was significant at trend-level, *t* = -1.86, *p* = .07.

**Table 1.** Fixed effects parameter estimates for multi-level model of efficacy ratings at baseline (1.1) and follow-up (1.2) as a function of strategy neurofeedback

| **1.1 Baseline Efficacy Ratings** |  |  |  |  | 95% *CI* |  |
| --- | --- | --- | --- | --- | --- | --- |
| Fixed effects (intercept, slopes) | Estimate | (SE) | *t* | *p* | Lower | Upper |
| Intercept | 6.05 | .21 | 8.8 | <.001 | 5.62 | 6.48 |
| Time | .79 | .20 | -3.9 | < .001 | -1.18 | -.39 |
| Between-subj, feedback | -.38 | .65 | -.58 | .56 | -1.66 | .90 |
| Between-subj, feedback x Time | -1.56 | .90 | -1.74 | .08 | -3.32 | .21 |
| Within-subj, feedback | -.06 | .62 | -.10 | .92 | -1.29 | 1.16 |
| Within-subj, feedback x Time | -1.64 | .88 | -1.86 | .07 | -3.38 | .10 |

| **1.2. Follow-Up Efficacy Ratings** |  |  |  |  | 95% *CI* |  |
| --- | --- | --- | --- | --- | --- | --- |
| Fixed effects (intercept, slopes) | Estimate | (SE) | *t* | *p* | Lower | Upper |
| Intercept | 5.26 | .22 | 4.4 | <.001 | 4.82 | 5.71 |
| Time | .786 | .20 | 3.9 | < .001 | .39 | 1.19 |
| Between-subj, feedback | -1.94 | .68 | 2.84 | < .01 | -3.28 | -.59 |
| Between-subj, feedback x Time | 1.56 | .90 | .74 | .08 | -.21 | 3.32 |
| Within-subj, feedback | -1.70 | .67 | 2.53 | .01 | -3.02 | -.37 |
| Within-subj, feedback by Time | 1.64 | .88 | 1.86 | .07 | -.10 | 3.38 |

*Note:* *N* = 13 persons, 2 timepoints, total of 208 observations. Due to the small sample size, we took a liberal approach to specifying degrees of freedom, based on the number of observations rather than the number of subjects. All *p*-values are two-tailed.

An identical model was run with frequency ratings as the dependent variable. None of the fixed-effect parameter estimates reached significance in the frequency model at baseline or follow-up (all p*s* > .1). A multi-level model with difficulty ratings was not run, given the lack of significant correlations between difficulty ratings and neurofeedback at baseline or follow-up.

## Discussion

The goal of the current study was to test a new technique for increasing metacognitive awareness and demonstrating the efficacy of previously-learned mood regulation skills in a sample of participants who had received cognitive behavioral therapy for depression. Our central hypothesis was that the memorable experience of using cognitive strategies in the scanner and receiving neurofeedback regarding the impact of those strategies on mood-related brain activation would affect participant beliefs and behavior. Specifically, we expected neurofeedback to increase the metacognitive awareness of the efficacy of specific strategies and thereby affect their likelihood of using those strategies in daily life. The significant correlations between neurofeedback presented in the scanner and the strategy ratings at follow-up, but not baseline, support this hypothesis: at both the between-subjects and within-subject level, stronger neurofeedback was associated with higher efficacy ratings four weeks after the scan. We predicted that the increased perceived efficacy would also translate into greater motivation to use the strategies; indeed, at the within-subject level, there was a marginally significant association between stronger neurofeedback and higher frequency of use four weeks after the scan

Two additional pieces of evidence support a causal relationship between the neurofeedback and post-scan changes in strategy ratings. First, effectiveness of a strategy in reducing activation in the target ROI after it had been activated by distressing thoughts and memories was not predicted by efficacy or frequency ratings obtained at the pre-scan baseline. Second, neurofeedback selectively affected follow-up ratings of experimental strategies used in the scanner; no relationship was observed between neurofeedback and ratings of unscanned control strategies. Importantly, experimental and control strategies did not differ in efficacy or frequency ratings at baseline, nor at follow-up; thus, the experience of using CBT strategies in the MRI session did not boost a general belief in the efficacy of all CBT strategies. Rather, only those strategies that were associated with neurofeedback evidence of their efficacy were rated as more effective and frequently used; moreover, these beliefs were still evident four weeks later. In sum, the findings that neurofeedback was associated with efficacy and frequency ratings only at follow-up—and only for strategies that worked to reduce brain activation in the target ROI—suggests that the context provided by the scan session made for a powerful learning experience with effects that generalized beyond the learning context itself.

Our approach diverges from most previous rt-fMRI training studies because we used neurofeedback as a tool to change beliefs about previously-learned skills rather than to teach or train a new skill. Time in the scanner allowed participants to test associations between their existing repertoire of strategies and changes in brain activation. To our knowledge, this study is the first to use rt-fMRI in this way as a tool for metacognitive awareness rather than skill training, and the first to provide participants who had received CBT with neurofeedback corresponding to their CBT strategy use. This novel approach has both theoretical and practical implications. Theoretically, it offers a salient example of therapeutic applications of declarative memory for enhancing metacognition. Pragmatically, it establishes a mechanistic paradigm for efficacious single-session neurofeedback interventions that could transform clinical practice.

### Developing a novel therapeutic approach

The current study represents a proof-of-concept for the development of a novel therapeutic intervention. It demonstrates the feasibility of providing ACC neurofeedback associated with the use of cognitive strategies learned in therapy for depression, or elsewhere. More broadly, it suggests a new avenue for rt-fMRI investigations that move beyond feedback-based training and take full advantage of a human participant’s ability to draw connections between an objective, biologically-based signal and subjective internal experience; the potential applications of such a metacognitive approach are numerous. While the current study establishes proof of principle and feasibility, future investigations will be required to demonstrate the efficacy and efficiency of this novel approach.

The small sample size in the current study—a consequence of our decision to recruit graduates of a clinical trial with standardized CBT experiences—is a significant limitation. Future iterations including a larger sample and independent control groups are now warranted. Control groups could receive no neurofeedback (as in Linden et al., 2012), or receive neurofeedback from a cortical region not ostensibly involved in mood regulation (as in Young et al., 2014). An additional limitation is that the results are primarily self-reported behavioral outcomes, so it is possible that the changes in self-reported efficacy and frequency ratings could reflect a reporting bias uncorrelated with the actual experience of use. Although the idea that metacognitions influence clinical course has significant supporting evidence, the relationship remains to be demonstrated in this application. Finally, during the pre-scan interview, participants were asked to list effective strategies for regulating their mood. The emphasis on identifying effective strategies for use in the study limited the range of efficacy ratings obtained at baseline (see Fig 4A). While this approach maximized participants’ chances of successful regulation in the scanner, it precluded our ability to observe neurofeedback-associated changes in strategies that were considered less effective/ineffective at baseline.

Potentially the most powerful application of this technique would be adjunct to a course of cognitive therapy. A majority of participants in the current study were no longer enrolled in therapy, and their memory of therapy content had likely degraded during the time since completing treatment. Providing patients with a neurofeedback experience *as strategies are being learned* in therapy could iteratively guide personalization of the therapy as well as motivate generalization of skill use outside of the session, accelerating treatment progress and improving long-term remission rates. Generalization of the protocol to different patient populations is also rational (Adcock et al., 2005), as the same mechanisms of change should apply transdiagnostically.

### Conclusions

Encouraged by these novel findings, we propose that a single-episode rt-fMRI intervention is a feasible and economical method to enhance the clinical impact of previously-learned cognitive strategies. By linking thoughts and mood states to brain activation, our participants gained 1) a mechanistic understanding of how mood is instantiated in the brain, 2) confidence in their ability to modulate their own brain activity, and 3) information about which strategies were most effective to guide future beliefs and motivation. Taken together, our findings offer the promise of a biologically-based, single-episode catalyst of behavior change that can be flexibly applied to enhance the generalization of therapy skills and improve clinical outcomes. We argue that rt-fMRI protocols such as the one tested here hold promise for helping patients understand the fundamental relationship between their subjective experience and biology. Indeed, this use of rt-fMRI technology to explore *auto-biology*—or self as a biological system (MacDuffie & Strauman, 2017)—has therapeutic implications for beliefs and behavior across a range of psychiatric disorders.

## Acknowledgements

The authors wish to thank Matthew Scult for his input on study design, Dr. Roger Beaty for his assistance with data collection, and Drs. Vishnu Murty and Shabnam Hakimi for their feedback on an earlier version of the manuscript. The study was supported by NIMH BRAINS award (R01MH094743) to R.A.A.

## References

Adcock, R. A., Lutomski, K., MacLeod, S., Soneji, D. J., Gabrieli, J. D., Glover, G., … deCharms, R. C. (2005). Real-Time fMRI During the Psychotherapy Session: Toward a Methodology to Augment Therapeutic Benefit, Exemplary Data. In Human Brain Mapping Annual Meeting. Toronto.

Deacon, B. J., & Baird, G. L. (2009). The chemical imbalance explanation of depression: reducing blame at what cost? Journal of Social and Clinical Psychology, 28(4), 415–435.

Eddington, K. M., Silvia, P. J., Foxworth, T. E., Hoet, A., & Kwapil, T. R. (2015). Motivational deficits differentially predict improvement in a randomized trial of self-system therapy for depression. Journal of Consulting and Clinical Psychology, 83(3), 602–616. https://doi.org/10.1037/a0039058

Fernandez-Duque, D., Evans, J., Christian, C., & Hodges, S. D. (2015). Superfluous Neuroscience Information Makes Explanations of Psychological Phenomena More Appealing. J Cogn Neurosci, 27(5), 926–944. https://doi.org/10.1162/jocn_a_00750

Hamilton, J. P., Glover, G. H., Hsu, J.-J., Johnson, R. F., & Gotlib, I. H. (2011). Modulation of subgenual anterior cingulate cortex activity with real-time neurofeedback. Hum Brain Mapp, 32(1), 22–31. https://doi.org/10.1002/hbm.20997

Hundt, N. E., Mignogna, J., Underhill, C., & Cully, J. A. (2013). The relationship between use of CBT skills and depression treatment outcome: A theoretical and methodological review of the literature. Behavior Therapy. Retrieved from http://www.sciencedirect.com/science/article/pii/S0005789412001177

Karlin, B. E., & Cross, G. (2014). From the laboratory to the therapy room: National dissemination and implementation of evidence-based psychotherapies in the U.S. Department of Veterans Affairs Health Care System. The American Psychologist, 69(1), 19–33. https://doi.org/10.1037/a0033888

Kazdin, A. E. (2007). Mediators and mechanisms of change in psychotherapy research. Annual Review of Clinical Psychology, 3, 1–27. https://doi.org/10.1146/annurev.clinpsy.3.022806.091432

Linden, D. E., Habes, I., Johnston, S. J., Linden, S., Tatineni, R., Subramanian, L., … Goebel, R. (2012). Real-time self-regulation of emotion networks in patients with depression. PLoS ONE, 7(6), e38115. https://doi.org/10.1371/journal.pone.0038115

MacDuffie, K. E., & Strauman, T. J. (2017). Understanding Our Own Biology: The Relevance of Auto-Biological Attributions for Mental Health. Clinical Psychology: Science and \ldots. Retrieved from http://onlinelibrary.wiley.com/doi/10.1111/cpsp.12188/full

Maldjian, J. A., Laurienti, P. J., Kraft, R. A., & Burdette, J. H. (2003). An automated method for neuroanatomic and cytoarchitectonic atlas-based interrogation of fMRI data sets. Neuroimage. Retrieved from http://www.sciencedirect.com/science/article/pii/S1053811903001691

McCabe, D. P., & Castel, A. D. (2008). Seeing is believing: The effect of brain images on judgments of scientific reasoning. Cognition, 107(1), 343–352.

Sulzer, J., Haller, S., Scharnowski, F., & Weiskopf, N. (2013). Real-time fMRI neurofeedback: progress and challenges. Neuroimage. Retrieved from http://www.sciencedirect.com/science/article/pii/S1053811913002759

Weisberg, D. S., Keil, F. C., Goodstein, J., Rawson, E., & Gray, J. R. (2008). The seductive allure of neuroscience explanations. Journal of Cognitive Neuroscience, 20(3), 470–477.

Young, K. D., Zotev, V., Phillips, R., Misaki, M., Yuan, H., Drevets, W. C., & Bodurka, J. (2014). Real-Time fMRI Neurofeedback Training of Amygdala Activity in Patients with Major Depressive Disorder. PLoS ONE, 9(2), e88785. https://doi.org/10.1371/journal.pone.0088785

